# Human brain organoid model of maternal immune activation identifies radial glia cells as selectively vulnerable

**DOI:** 10.1101/2022.08.09.503336

**Authors:** Kseniia Sarieva, Theresa Kagermeier, Shokoufeh Khakipoor, Ezgi Atay, Zeynep Yentür, Katharina Becker, Simone Mayer

## Abstract

Maternal immune activation (MIA) during the critical windows of gestation is correlated with long-term neurodevelopmental deficits in the offspring, including increased risks for autism spectrum disorder (ASD) in humans. Interleukin 6 (IL-6) derived from the gestational parent is one of the major molecular mediators, by which MIA alters the developing brain. In this study, we established a human three-dimensional (3D) *in vitro* model of MIA by treating induced pluripotent stem cell-derived dorsal forebrain organoids with a constitutively active form of IL-6, Hyper-IL-6. We validated our model by showing that dorsal forebrain organoids express the molecular machinery necessary for responding to Hyper-IL-6 and activate STAT signaling upon Hyper-IL-6 treatment. RNA sequencing analysis revealed the upregulation of major histocompatibility complex class I (MHCI) genes, which have been implicated with ASD. Immunohistochemical analysis as well as single-cell RNA-sequencing revealed a small increase in the proportion of radial glia cells. Single-cell transcriptomic analysis revealed the highest number of differentially expressed genes in radial glia cells with downregulation of genes related to protein translation in line with data from mouse models of MIA. Additionally, we identified differentially expressed genes not found in mouse models of MIA which might drive species-specific responses to MIA. Together, we establish a human 3D model of MIA, which can be used to study the cellular and molecular mechanisms underlying the increased risk for developing disorders such as ASD.

## Introduction

The role of environmental factors in shaping prenatal brain development and thus contributing to neurodevelopmental disorders has been increasingly appreciated over the last years (1). Epidemiological studies suggest that health of gestational parent (i.e. women in the experimental setup) correlates with fetal neurodevelopmental outcomes (2–4). For example, infections of the gestational parent have been associated with long-lasting neurodevelopmental deficits in offspring, including autism spectrum disorder (ASD) (5, 6). The data collected from the offspring of gestational parents who were admitted to hospital due to infection suggest an ASD hazard ratio of 2.98 (95% CI 1.24-7.15) due to viral infection in the first trimester and 1.42 (95% CI 1.08–1.87) for bacterial infection in the second trimester (5). Especially with a large number of Covid-19 cases in pregnant people in the last years, it will be important to consider possible effects of maternal Covid-19 infection on the offspring in the near future (7). Common to different types of infection is the increase in cytokines and chemokines that, either via placental transfer or induction of placental inflammation and subsequent cytokine and chemokine release, leads to elevated concentrations of these immune mediators in the fetal compartment with consequences for the developing brain (7). Elevated concentrations of the pro-inflammatory cytokine interleukin-6 (IL-6) during pregnancy have been associated with newborn offspring brain anatomy as well as structural and functional connectivity with consequences for cognitive function in infancy (8–10). Taken together, while there is mounting evidence for maternal immune activation (MIA) to affect neurodevelopmental trajectories and psychiatric disease risk in humans, we currently do not understand the molecular mechanisms that translate infections of the gestational parent to adverse neurodevelopmental outcomes.

Diverse animal models have shown that MIA contributes to the behavioral deficits that recapitulate ASD symptoms in humans (11–15). In these models, MIA is generally modeled by mimicking viral infection through a single injection of synthetic double-stranded RNA (dsRNA), polyinosinic:polycytidylic acid (poly(I:C)) at E12.5-18 in mice (16, 17). Using gain and loss-of-function approaches in such models, several cytokines including IL-17α and IL-6 have been identified to be causal for behavioral deficits (11, 16, 17). Mechanistically, IL-17α is suggested to mediate abnormal protein translation specifically in the neurons of male offspring resulting in behavioral deficits reminiscent of ASD (16). IL-6 has been shown to be necessary and sufficient to induce behavioral abnormalities in male offspring (11). Additionally, MIA induces an increase in the proliferation of radial glia cells (RGs) in the neocortex in an IL-6-dependent manner in mice (18). This resulted in an overproduction of deep-layer excitatory neurons later during development (18). Together, rodent studies provide valuable insights into the molecular mechanisms of MIA suggesting ASD-relevant changes in both neural progenitor cells and neurons in the developing brain (11, 16–19). However, these studies suffer from limitations including within-litter variability in behavioral responses (20) and a lack of translational potential to humans due to interspecies differences (21).

Instead, recent progress in human- *in vitro* models now allows investigating MIA in a human cellular context. In line with this idea, a recent study used human induced pluripotent stem cells (iPSCs)- derived neural precursor cells (NPCs) to study how interferon-γ (IFN-γ) may mediate some effects of MIA (22). In this model, IFN-γ activates the anti-viral response resulting in up-regulation of MHCI genes, targeting MHCI proteins to the growth cone and, consequently, abnormal neurite outgrowth (22). This effect is accompanied by ASD-relevant transcriptional abnormalities (22).

Recent advances in PSCs-derived 3D brain organoid cultures provide new avenues to investigate the effects of environmental adversities on early brain development in a human context (21). Brain organoids model key aspects of the cellular complexity of the developing human brain *in vitro* thus enabling direct observations of the processes normally inaccessible *in utero* (23). Together with single-cell omics technologies, we propose that brain organoids may be a powerful tool for studying cell type-specific effects of molecular mediators of MIA, including IL-6.

IL-6 acts on its target cells by binding to its specific receptor, interleukin-6 receptor (IL6R), which forms a complex with interleukin-6 signaling transducer (IL6ST), also known as gp130 (Figure 1a) (24). In classic signaling, IL6R and IL6ST are expressed by the same cell in *cis* (24). However, Il6R can also be supplied in *trans* as a soluble form (s-IL6R) originating either from the alternative splicing of IL6R gene or from shedding from plasma membrane by metalloproteinases (24, 25). Interestingly, a recent preprint found that human iPSC-derived microglia are constitutively releasing s-IL6R (26). In absence of microglia, t*rans* signaling can be modelled by exposing target cells to the chimeric protein Hyper-IL-6 in which s-IL6R is covalently bound to IL-6 through a flexible peptide link (Figure 1b) (27). Upon binding of Hyper-IL6 to IL6ST, intracellularly, IL6ST is autophosphorylated leading to the activation of the Janus kinase/signal transducer and activator of transcription (JAK/STAT) pathway (24) (Figure 1c).

**Figure 1.**
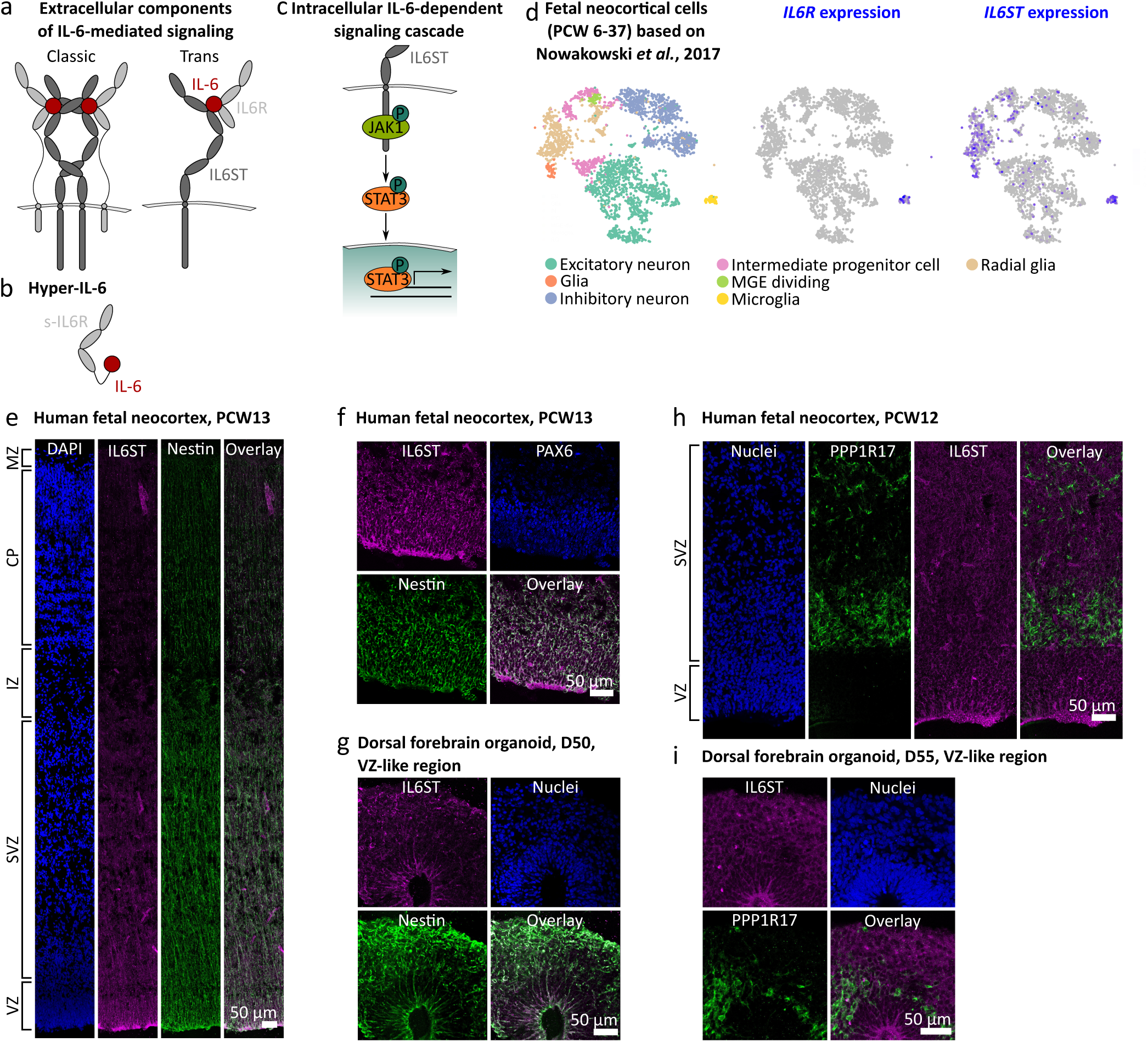
Characterization of the IL-6 signaling pathway in human midgestational neocortex and dorsal forebrain organoids. a, IL-6 has two receptors, IL6R and IL6ST. When both of them are expressed in the target cell, IL-6-dependent signaling cascade is called “classic” whereas when IL6R is provided extracellularly for example by a different cell in its soluble form, the cascade is called “trans”. b, Hyper-IL-6 is a chimeric protein consisting of soluble IL6R (s-IL6R) and IL-6 connected to each other through a peptide linker. c, Upon binding to IL6R and IL-6, IL6ST induces phosphorylation of JAK1, which in turn phosphorylates STAT3 at Y705. Then, p-Y705-STAT3 enters nucleus and activates transcription. d, In the developing human neocortex, IL6R is expressed exclusively in the microglia whereas IL6ST expression can be detected in radial glia and to a lower extend in intermediate progenitor cells, as well as in glial cells in a single-cell RNA-sequencing dataset. e, IL6ST (magenta) immunofluorescence of human PCW13 neocortex, shows protein expression in the VZ and SVZ. Nestin (green) is used as a marker of radial glia cells. VZ, ventricular zone; SVZ, subventricular zone; IZ, intermediate zone; CP, cortical plate; MZ, marginal zone. f, In the VZ, IL6ST immunofluorescence overlaps with Nestin at PCW13. g, IL6ST immunofluorescence is associated with Nestin in the VZ-like area of dorsal forebrain organoids at day 50 of differentiation. h, IL6ST (magenta) in human neocortex at PCW12 does not co-localize with PPP1R17 (green) used as a marker of intermediate progenitor cells. I, IL6ST immunofluorescence is not associated with PPP1R17 in the dorsal forebrain organoids at day 55 of differentiation.

The IL6ST/JAK/STAT signaling pathway plays an important role in the developing of neocortex in physiological conditions, a brain region predominantly affected in many neurodevelopmental disorders including ASD (28). IL6ST acts as an accessory receptor to other cytokines of the IL-6 family, including leukemia inhibitory factor (LIF), which contributes to maintaining the radial glial pool and is partially supplied by the cerebrospinal fluid (29, 30). The IL6ST-dependent signaling pathway is activated in some SOX2-positive RGs in the germinal zones of the developing human neocortex at midgestation (31). However, IL-6 itself is hardly expressed in the developing human and mouse brain (32, 33), and a rodent radioisotope tracing study suggests that IL-6 in the prenatal brain is primarily of maternal origin (34). Therefore, maternal IL-6 signaling through JAK/STAT may be an important mediator of MIA in the developing human brain.

In this study, we developed a human 3D *in vitro* model of MIA using dorsal forebrain organoids (DFOs) (35) and activating IL-6 signaling to reveal the cellular and molecular responses to MIA during neocortical development. Since we found IL6ST expression in RGs in DFOs recapitulating expression in primary human tissue, we used IL-6/Hyper-IL-6 exposure for 5 to 10 days in DFOs starting at day 45 of differentiation to model MIA. This time point corresponds to the human neocortex at early midgestation. We treated cells for several days to account for extended period of elevated cytokine levels upon exposure to infections. We validate our model by showing activation of JAK/STAT signaling upon Hyper-IL-6 exposure as well as upregulation of transcripts encoding proteins involved in the innate immune response. Immunohistochemical analysis of cell type composition revealed minor effects of Hyper-IL-6 treatment on the number of SOX2-positive ventricular RGs (vRGs). Using single-cell RNA sequencing (scRNAseq) and trajectory analysis, we found that exposure to Hyper-IL-6 led to a shift towards earlier progenitor cell types at the expense of neurons in the excitatory lineage. Hyper-IL-6 induced cell type-specific transcriptional changes, including upregulation of inflammation-related genes and downregulation of genes involved in protein translation specifically in vRGs in agreement with previous animal and *in vitro* models of MIA. Moreover, our 3D model of MIA also revealed differentially expressed genes not reported in any other model, which could have species-specific effects and may be especially relevant to link to human epidemiological findings. We found NR2F1, an important regulator of brain development (36), to be upregulated across cell clusters. Therefore, our model is a needed extension to previous models of MIA and opens the door for further investigating causal relations underlying MIA-dependent neurodevelopmental trajectories.

## Materials and Methods

### iPS cell culture and dorsal forebrain organoids differentiation

Human iPSC lines BIONi010-C (Source: EBiSC) and HMGU1 (Source: Human Pluripotent Stem Cell Registry) were cultured in standard conditions (37°C, 5% CO_2_, and 100% humidity), in E8 flex medium (Gibco, Cat. no. A2858501) and passaged in colonies using Gentle Dissociation Reagent (STEMCELL Technologies, Cat. no. 07174) onto hESC-qualified growth factor-reduced Matrigel-coated (Corning, Cat. no. 354277) plates. Cell lines were maintained below passage 25, and were routinely tested for mycoplasma contamination (assayed with PCR Mycoplasma Detection Set, TaKaRa, Cat. no. 6601) and pluripotency status with antibodies against OCT4 (rabbit, 1:500, Abcam, Cat. no. ab19857) upon thawing each cryovial.

Dorsal forebrain organoids were generated as previously described with minor modifications (35). Briefly, iPSCs at ∼70-80% confluency were dissociated into single cells using Accutase (MERCK, Cat. no. A6964) and plated at 9 000 cells per well in 96 well V-bottom low adhesion plates (S-bio, Cat. no. MS-9096VZ) in cortical differentiation medium I (CDMI) supplemented with 20 µM Y-27632 (Cayman Chemical, Cat. no. 10005583), 5 µM SB 431542 (Tocris, Cat. no. 1614), and 3 µM IWR-1 (MERCK, Cat. no. 681669). Glasgow’s MEM (GMEM)-based (Gibco, Cat. no. 11710035) CDMI includes 20% KnockOut Serum Replacement (KOSR, Thermofisher Scientific, Cat. no. 10828028), 1X non-essential amino acids (Sigma, Cat. no. M7145), 0.11 mg/mL Sodium Pyruvate (Thermofisher Scientific, Cat. no. 11360070), 1X Penicillin-Streptomycin (Thermofisher Scientific, Cat. no. 15140122), 0.1 mM β- Mercaptoethanol (Gibco, Cat. no. 21985023). At this stage, media was changed every three days. After six days of differentiation, Y-27632 was withdrawn by media change. After 18 days, dorsal forebrain organoids were transferred from 96- to 24-well low adhesion plates and moved to an orbital shaker rotating at 53 rpm (2.5 cm throw) with media exchange to CDMII (DMEM/F12-based medium containing 1X Glutamax (Thermofisher Scientific, Cat. no. 31331093), 1X N2 Supplement (Thermofisher Scientific, Cat. no. 17502048), 1X CD Lipid Concentrate (Thermofisher Scientific, Cat. no. 11905031) and 1X Penicillin-Streptomycin (Thermofisher Scientific, Cat. no. 15140122)). At 35 days, organoids were moved into CDMIII (DMEM/F12-based medium containing 10% FBS (GE Healthcare Life Sciences, Cat. no. SH30070.03), 5 µg/mL Heparin (MERCK, Cat. no. H3149-25KU), 1X N2 Supplement, 1X CD Lipid Concentrate and 1% Matrigel (Corning, Cat. no. 356234)). At 70 days, CDMIV was introduced by additionally supplementing CDMIII with 1X B27 Supplement (Thermofisher Scientific, Cat. no. 17504044). Starting from day 18, media changes were performed every 3-4 days.

To treat organoids with IL-6 and Hyper-IL-6, the compounds were diluted in 0.1% BSA (AppliChem, Cat. no. A0850) in PBS (also used as vehicle control). At the start of treatment, a complete media change was performed with CDMIII containing respective compounds at a final concentration of 8.8 ng/ml (422.6 nM) IL-6 (R&D Systems, Cat. no. 7270-IL-025) and 25 ng/ml (422.6 nM) Hyper-IL-6 (R&D Systems, Cat. no. 8954-SR-025) at day 45 of differentiation. The treatment was maintained by daily half-volume media change until termination of the experiment or treatment withdrawal at day 55 of differentiation.

### Human tissue samples

De-identified second trimester tissue samples were collected with previous patient consent in strict observance of the legal and institutional ethical regulations. Protocols were approved by the Human Gamete, Embryo, and Stem Cell Research Committee (Institutional review board) at the University of California, San Francisco and the University Hospital Tübingen. Tissues were fixed, cryopreserved and cryosectioned at 16 µm as described in (37).

### Immunohistochemistry in organoid slices and human tissue samples

Whole dorsal forebrain organoids were fixed in 4% paraformaldehyde (PFA) in PBS for 45–60 min at room temperature (38). Organoids were washed three times with PBS and then incubated in 30% sucrose in PBS solution until saturated. Organoids were embedded in a 1:1 v:v mixture of 30% sucrose in PBS and optimal cutting temperature (OCT) compound (TissueTek) and sectioned on Superfrost Plus slides (VWR) with a cryostat at 20 µm (Leica).

For immunostaining, slides were thawed to room temperature, washed with PBS before permeabilization and blocking with 1% Triton-X100, 0.2% gelatin, and 10% normal donkey serum in PBS for 1 hour. The following primary antibodies were used: anti-Ki-67 (rabbit, 1:300, Merck, Cat. no. AB9260), anti-SOX2 (goat, 1:500, R&D Systems, Cat. no. AF2018), anti-PAX6 (mouse, 1:300, Atlas Antibodies, Cat. no. AMAb91372), anti-CTIP2 (rat, 1:400, Abcam, Cat. no. ab18465), anti-TBR2 (rabbit, 1:100, Atlas Antibodies, HPA028896), anti-SATB2 (rabbit, 1:500, Abcam, ab34735). For immunostainings with anti-TBR2 and anti-PAX6 antibodies, antigen retrieval in boiling 10 mM citric acid buffer (pH 6.0) was performed. Primary antibodies diluted in permeabilization and blocking solution were applied to the sections overnight at 4 °C. After washing three times with PBS, secondary antibodies diluted in permeabilization and blocking solution were applied to the sections for 3 hours at room temperature. Finally, sections were washed three times with PBS and stained with the nuclear dyes DAPI (Thermofisher Scientific, Cat. no. D1306) or DR (Thermofisher Scientific, Cat. no. 62251) following the respective protocols supplied by the manufacturer. Images were acquired by Leica SP8 confocal microscope at 40x magnification objective. Quantitative analyses were conducted on three or more randomly selected rosette structures per section of an organoid unless stated otherwise. The numbers of cells positive for each marker were measured with Imaris 9.7 software using “Surfaces” and “Spots” options (Bitplane, RRID:SCR_007370). To quantify the area, perimeter and number of ventricular zone (VZ)-like regions, the VZ was defined by dense SOX2+ immunoreactivity. To quantify the distribution of CTIP2+/SATB2+ cells, the cell numbers were quantified in 200 µm-wide regions of interest starting from the outer surface of the organoid (three randomly chosen positions on the section of the organoid).

### Western Blotting

Organoids were lysed in 1X RIPA buffer (Cell Signaling technology, Cat. no. 9806) in the presence of 1x Halt protease and phosphatase inhibitor cocktail (Thermofisher Scientific, Cat. no. 78443) (39). Subsequently, the samples were incubated on rotating wheel for 1 hour, sonicated using probe sonicator (20 seconds, 10 pulses, 50% power, Bandelin, SONOPULS mini20, MS1.5 probe) and centrifuged at 14 000 g for 30 minutes at 4°C. The total protein concentrations were determined using the Pierce™ BCA assay (Thermofisher Scientific, Cat. no. 23225). Samples containing equal amounts of protein (5-10 μg) were boiled in complete sample buffer (Li-COR, Cat. no. 928-40004) at either 95°C for 5 minutes or at 70°C for 10 minutes and resolved in a 4-15% Precast Gels (Bio-Rad, 12-well, Cat. no. 456-1095, 15-well, Cat. no. 456-1086). Proteins were transferred onto Immun-Blot Low Fluorescence PVDF membrane (Bio-Rad, Cat. no. 1620264) using Trans-blot Turbo semi-dry transfer system. Total protein staining in membranes was performed using Revert™ 700 Total Protein Stain Kit for Western Blot Normalization (Li-COR, Cat. no. 926-11016). The membranes were blocked with Tris-buffered saline-based blocking buffer (Li-COR, Cat. no. 927-60001) for 1 hour at room temperature. Subsequently, the membranes were incubated with the primary antibodies diluted in Tris-buffered saline-based antibody dilution buffer (Li-COR, Cat. no. 927-65001) at 4 °C overnight. The following primary antibodies were used: anti-STAT3 (mouse, 1:1000, Cell Signaling Technologies, Cat. no. 9139), anti-phospho-Y705-STAT3 (rabbit, 1:1000, Cell Signaling Technologies, Cat. no. 9145), anti-NR2F1 (rabbit, 1:1000, Abcam, Cat. no. ab181137) and anti-β-actin (mouse, 1:5000, Abcam, Cat. no. ab6276). After washing in 1X TBS-T (Tris-buffered saline pH 7.5, 0.1% Tween-20), the membranes were incubated with anti-rabbit (Li-COR, Cat. no. 926-32211) and anti-mouse (Li-COR, Cat. no. 926-32211) secondary antibody conjugated with infra-red dyes for 1 hour at room temperature. Blots were then imaged with Odyssey Scanner and analyzed using Image Studio and Image Studio Lite v.5 software (Li-COR) with normalization to the background and Total Protein Stain.

### RNA sequencing

mRNA was isolated from organoids using the RNeasy Mini Kit (Qiagen, Cat. no. 74104) and RNase-Free DNase set (Qiagen, Cat. no. 79254). Library preparation was conducted by Novogene (https://en.novogene.com/) using non-directional poly-A enrichment strategy. Samples were then sequenced on the Illumina NovaSeq 6000 platform using paired-end, 150-bp-long reads to 30 million reads per sample. HISAT2 was used to align raw reads to the GRCh37/hg19 human genome reference (40). Reads containing adaptors, more than 10% of undefined bases and more than 50% of bases with Qscore (Quality value) <= 5 were excluded from further analysis. Gene expression levels were estimated using FPKM (expected number of Fragments Per Kilobase of transcript sequence per Millions base pairs sequenced) (41). Principal component analysis (PCA) was performed on the FPKM values. Hierarchical clustering was used to cluster the FPKM values of genes. Differential gene expression (DGE) analysis was performed using the DESeq2 R package (1.10.1) (42). P-values were adjusted for multiple comparisons using the Benjamini-Hochberg method and significantly differentially expressed genes were selected based on two criteria: FDR ≤ 0.2 and absolute log2 Fold Change > 0.4 between the Vehicle and Hyper-IL-6 conditions. Functional analysis of the differentially expressed genes was performed using clusterProfiler (43). Weighted gene co-expression analysis (WGCNA) was performed using normalized FPKM values. The WGCNA R package computed an unsigned similarity matrix with a softpower of 10 (44). Gene modules were created on the basis of hierarchical clustering results of the similarity matrix and eigengene per module was calculated. The relationship between gene modules and experimental conditions was evaluated by comparing eigengene values by two-sided t-test with a Bonferroni correction for multiple comparisons. The majority of bioinformatics analysis was conducted by Novogene (https://en.novogene.com/).

### Single-cell dissociation and multiplexing for single-cell RNA sequencing

Individual dorsal forebrain organoids were dissociated using the Worthington Papain Dissociation System kit (Worthington Biochemical, Cat. no. LK003150) following a published protocol (45) with minor modifications. Organoids were cut with a scalpel blade and then placed in 2.5 ml papain supplemented with DNAse inhibitor and incubated at 37 °C for 15 min while shaking at 57 rpm (2.5 cm throw). Organoids were gently triturated ten times using a 10-ml serological pipette and a 1-ml pipette, and incubation continued for another 10 min. Next, organoids were triturated ten times using a 10-ml serological pipette. Large debris was allowed to settle down in the Ovomucoid Inhibitor diluted in Earle’s medium, and the supernatant containing single cells was centrifuged at 300xg for 7 minutes at room temperature. Cells were resuspended in 0.04% BSA in PBS and passed through a 40-µm cell strainer (Flowmi). Dead Cell Removal (Miltenyi, Cat. no. 130-090-101) was performed following the manufacturer’s instructions. After final centrifugation, the cells were subjected to multiplexing using 3’ CellPlex Kit Set A (10x Genomics, Cat. no. PN-1000261) following the manufacturer’s instructions. Finally, the cells were quantified and pooled in equal proportions. Approximately 16 500 cells per channel were loaded onto a Chromium Single Cell 3′ Chip (10x Genomics, Cat. no. PN-120236) and processed through the Chromium controller to generate single-cell gel beads in emulsion. scRNAseq libraries were prepared with the Chromium Single Cell 3′ Library & Gel Bead Kit v.3 (10x Genomics, Cat. no. PN-1000121). Libraries from different samples were pooled and sequenced on a NextSeq 500 instrument (Illumina).

### Single-cell RNA sequencing analysis

#### Quality control and preprocessing

scRNAseq data were aligned to the GRCh38-0.1.2 reference genome, and single-cell gene read count tables were generated using the CellRanger (v.3.6.0) pipelines multi and aggr without normalization. Data were analyzed in R (v4.0) using Seurat (v4.0.4) (46). Quality control removed cells with fewer than 1 000 and more than 7 500 genes per cell, cells with log10(genes per UMI) less than 0.8 and cells with greater than 25% mitochondrial content. Gene expression was normalized using the SCTransform workflow and the different samples were then integrated using the integration workflow as published (47). Dimensionality reduction was performed using PCA, and we selected 30 PCs based on the Elbow plot. Louvain clustering was performed at resolutions from 0.2 to 1.0, and resolution 0.4 was chosen based on clustree cluster stability assessment (48). UMAP plots were generated with Seurat package default parameters. Doublets were removed using DoubletFinder (49). For identifying the markers for each cluster, the FindAllMarkers function was used with both MAST (50) and roc algorithms applied to the raw count matrix. Additionally, we manually assessed expression of canonical genes to assign each cluster to a known cell type. Brain regional identity of individual clusters was assessed using the VoxHunt algorithm and SCTransform expression values were used as input (51).

#### Cell proportion analysis

Differences in cell type proportions between experimental conditions were analyzed with a permutation test followed by bootstrapping (https://github.com/rpolicastro/scProportionTest) where clusters with FDR less than 0.05 and absolute log2 fold change more than 0.58 were considered differentially abandoned.

#### Trajectory inference and trajectory quality assessment

Slingshot algorithm was applied to construct putative single-cell differentiation trajectories based on the UMAP representation (52). Pseudotime zero was assigned to the vRG cell cluster. A random forest classifier was used to retrieve genes differentially expressed along the putative deep-layer neuron differentiation trajectory.

#### Identification of differentially expressed genes, gene module signatures and gene set enrichment analysis

To identify differentially expressed genes between experimental conditions by cell type, we performed a DGE analysis using DESeq2 as implemented in the Seurat package. To control for differences in cluster representation between the samples, a random selection of N cells per cluster was chosen where N was the minimal cell count for each cluster across samples. Genes with FDR < 0.05 were considered statistically significant. Gene set enrichment analysis was performed using the clusterProfiler R package (FDR < 0.05 for significant gene ontology terms) (43). DEGs were annotated as transcription factors based on (53).

To calculate differential expression of modules, SCTransform normalized count matrix was supplied to UCell algorithm using a predefined gene list from selected GO terms (54).

Additional gene set enrichment analysis was performed for ASD risk genes from the SFARI database (categories 1-4 from SFARI release on 31.10.2019; categories 1-2 from SFARI release on 20.07.2022) (55). Enrichment was tested using a one-sided Fisher’s exact test in R to test whether the proportion of risk genes in the differentially expressed set in each cell cluster is more than expected by chance. Genes expressed in our dataset as assessed by DESeq2 were used as background. Up- and down- regulated genes were tested simultaneously. P-values were corrected for multiple tests (by number of cell clusters) using the Benjamini-Hochberg method.

#### SCENIC analysis

A single-cell regulatory network for cycling vRGs was constructed with SCENIC (56). Specifically, GENIE3 was applied to infer gene regulatory networks from log-normalized count data (57). For identification and scoring gene regulatory networks, or regulons, AUC was calculated using AUCell. Regulon specificity scores were calculated based on (58).

### Statistics

The data distribution was not assumed to be normal in most cases and, therefore, Aligned Rank Transform (ART) was used for statistical testing unless stated otherwise. The cell line of origin was included as a covariate in the models. Organoids were randomized into different treatment groups with approximately equal numbers of organoids in each group whenever possible. Details of specific statistical comparisons are listed in the relevant figure legends.

## Results

### Radial glia in the human neocortex at midgestation express the molecular machinery to respond to IL-6 exposure

To determine cell types susceptible to IL-6 exposure in the developing human neocortex at midgestation, a peak period for neurogenesis, we first analyzed the expression of the IL-6 receptors, IL6R and IL6ST, in a scRNAseq dataset generated by Nowakowski and colleagues (32). We found that IL6R was expressed exclusively in microglia (Figure 1d), whereas IL6ST was predominantly present in both microglia and RGs (Figure 1d). We confirmed IL6ST expression in the ventricular (VZ) and subventricular (SVZ) zones *in situ* (Figure 1e) and found association with the RG marker Nestin (Figure 1f) but not with the marker of intermediate progenitor cells (IPCs) PPP1R17 (Figure 1h and Supplementary Figure 1a). As reported previously (31), we found that active STAT (p-Y705-STAT3) co-localized with SOX2, a nuclear marker of RGs, in a subset of cells in both VZ and SVZ (Supplementary Figure 1b). We therefore concluded that human radial glia cells as midgestation might be susceptible to IL-6 through *trans* signaling (Figure 1a) with IL6R supplied by microglia.

Similar to the human neocortex at early midgestation, human DFOs generated using a previously published protocol (35) at day 50 of differentiation express IL6ST in the Nestin-positive RGs localized to the VZ-like area (Figure 1g) and did not show co-localization of IL6ST with PPP1R17 (Figure 1i). At this stage, organoids consist mainly of SOX2-positive RG-like cells organized into rosettes, TBR2- positive IPCs, and CTIP2-positive deep-layer excitatory neurons (Supplementary Figure 1c). This cellular arrangement corresponds to the human neocortex at early midgestation (Supplementary Figure 1d) (32), but lacks microglia.

### Establishment and characterization of IL-6 treatment with human dorsal forebrain organoids

To investigate the effect of IL-6 exposure on human neocortical development, we differentiated human DFOs from two male iPSCs lines since pronounced effects are expected to be found in males based on animals studies (16) and ASD prevalence in humans (59). We exposed DFOs to 8.8 ng/ml (422.6 nM) IL-6 and equimolar concentration of Hyper-IL-6 (25 ng/ml, 422.6 nM) for 5 to 10 days starting at day 45 of differentiation to model the period of early midgestation. This concentration is comparable with the serum concentration of IL-6 upon septic infection (60). The prolonged treatment scheme was chosen because IL-6 serum concentrations can be elevated beyond acute inflammatory response and time points were analyzed independently unless stated otherwise (61). 0.1% BSA diluted in PBS was used as a vehicle solution for both IL-6 and Hyper-IL-6 to enhance the stability of recombinant proteins (see Figure 2a for experimental design). To assess the activation of the signaling cascade downstream to IL-6/Hyper-IL-6 treatment, we analyzed phosphorylation of STAT3 at Y-705 relative to total STAT3 (Figure 2b). We found that Hyper-IL-6 but not IL-6 resulted in an increased p-Y705-STAT3 (Figure 2c). We, therefore, concluded that, as predicted by gene expression data (Figure 1), DFOs were capable of responding to Hyper-IL-6 treatment through *trans* signaling. Based on this finding, we reasoned that Hyper-IL-6 treatment could be used to model MIA in DFOs.

**Figure 2.**
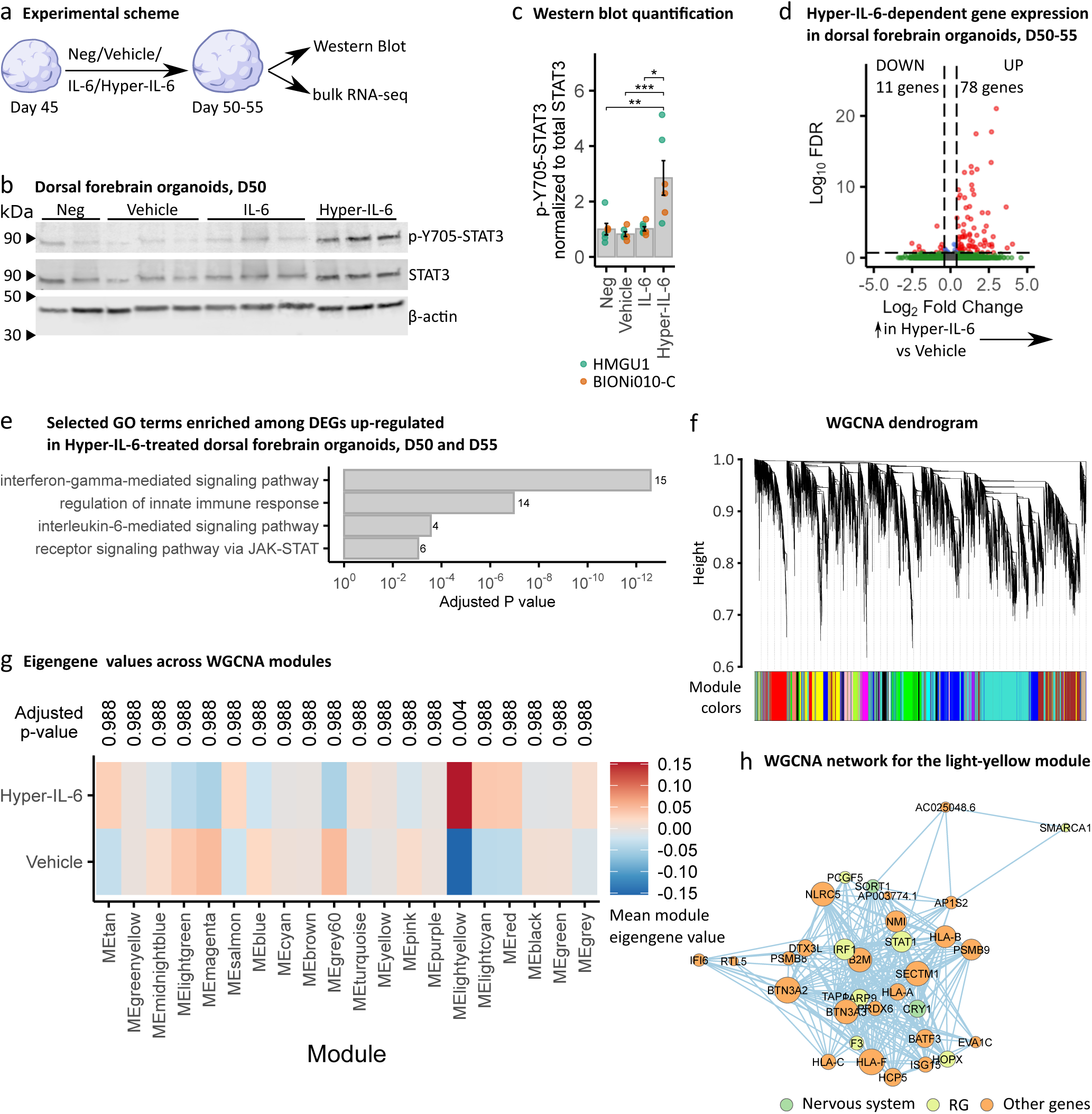
Hyper-IL-6 treatment leads to the activation of JAK/STAT intracellular cascade and results in transcriptional changes in dorsal forebrain organoids. a, Scheme of the cytokine treatment in dorsal forebrain organoids. Neg, negative control; Vehicle, vehicle control (0.1% BSA in PBS); IL-6, 8.8 ng/ml in 0.1% BSA in PBS; Hyper-IL-6, 25 ng/ml in 0.1% BSA in PBS. b, Representative Western Blot of phospho-Y707-STAT3 (top), STAT3 (middle) and β-actin (bottom) in individual dorsal forebrain organoids at day 50 of differentiation after 5 days of treatment. c, Quantification of signal intensity of phospho-Y705-STAT3 relative to total STAT3 from b. Each dot represents an individual organoid sample. Color represents cell line of origin, HMGU1 (green), BIONi010-C (orange). Neg (n=6), Vehicle (n = 6), IL-6 (n=7) and Hyper-IL-6 (n = 6) DFOs from one batch per iPSC line. Bars represent mean, error bars represent ± SEM. Comparisons were analyzed using Aligned Rank Transform (ART) ANOVA: *, p-value < 0.05, **, p-value < 0.01; ***, p-value < 0.001. d, Volcano plot of Hyper-IL-6- dependent gene expression in dorsal forebrain organoids at days 50 and 55 of differentiation treated for 5 to 10 days. Red dots indicate statistical significance (FDR < 0.2, absolute log2 Fold Change > 0.4). Positive log2 Fold Change indicates higher gene expression in Hyper-IL-6-treated relative to Vehicle-treated dorsal forebrain organoids. Data from n=12 organoids per condition (two cell lines, day 50 and 55 analyzed together). e, GO overrepresentation analysis of differentially upregulated genes (FDR < 0.2, log2 Fold Change > 0.4) between Hyper-IL-6 and Vehicle-treated dorsal forebrain organoids. The x axis displays the adjusted p-value. Numbers next to the bars represent the number of the differentially expressed genes belonging to the GO term. Data from n=12 organoids per condition (two cell lines, day 50 and 55 analyzed together). f, Dendrogram for WGCNA obtained by clustering the dissimilarity based on consensus Topological Overlap. Each color represents a module, which contains a group of highly connected genes. A total of 20 modules were identified. g, Heatmap representing mean module eigengene value by treatment condition based on WGCNA. Comparisons were analyzed using t-test, Bonferroni adjusted p-value. h, WGCNA network for the light-yellow module. Color of the dots represents gene group: nervous system-enriched, green; radial glia-enriched, yellow; other genes including innate immune response genes, orange.

STAT3 is a transcriptional activator previously reported to have cell type-specific targets in human neurodevelopment (31, 62, 63). Moreover, MIA has previously been shown to induce transcriptional changes in rodent models (16, 64). Therefore, we next investigated the transcriptional changes associated with the exposure of DFOs to Hyper-IL-6 by performing RNAseq on days 50 and 55 of differentiation. To identify gene expression changes driven by Hyper-IL-6, we performed differential gene expression (DGE) analysis irrespective of the length of treatment. In total, we identified 89 differentially expressed genes (DEGs) (Figure 2D; FDR ≤ 0.2, absolute log2 Fold Change ≥ 0.4); three DFOs from two iPSC lines in two differentiation experiments) of which only 11 genes showed a downregulation. The upregulated DEGs were associated with innate immune response (e.g. MHCI members and B2M) as well as with response to IL-6 and signaling through the JAK/STAT pathway (Figure 2e and Supplementary Table 2). Interestingly, MHCI components and its accessory B2M protein have been reported to be also deregulated upon IFN-γ treatment in human iPSC-derived NPCs (22).

Since gene expression has a correlative structure it can be analyzed as gene modules (65). These gene co-expression modules enable functional interpretation of the DGE relevant for neurodevelopmental diseases (66). We therefore employed weighted gene co-expression network analysis (WGCNA) to gain insights into potential correlative gene expression characterizing Hyper-IL- 6 treated organoids (Figure 2f). WGCNA revealed 16 modules of genes with correlated expression. Only the light-yellow module was associated with Hyper-IL-6 treatment (Figure 2g). A network connection representation of the genes constituting the light-yellow module showed an overlapping expression of RG-enriched genes with genes involved in inflammatory response (Figure 2h). This result suggests that among the cell types that are present in the DFOs, RGs are the most susceptible to Hyper-IL-6 exposure.

### Cell type composition of the DFOs is largely undisturbed by the Hyper-IL-6 treatment

Our findings suggest that Hyper-IL-6 treatment is mostly targeting RGs within DFOs. Previous studies have indicated that MIA in mice alters the proportions of proliferative versus neurogenic cell divisions at E12.5 culminating in an overproduction of deep-layer neurons later in development (18). To evaluate potential disturbances in the cell fate decisions of RGs upon exposure to Hyper-IL-6, we performed a series of immunohistochemical analyses to assess organoid morphology and the cell type compositional changes (Figure 3a). First, we followed the growth rate of the DFOs upon Hyper-IL-6 treatment on days 45-55 and until day 90 of differentiation. We observed no differences in the organoid size (Figure 3b) nor in area, perimeter and number of SOX2-positive VZ-like regions within individual organoids at days 50 and 55 of differentiation (Figure 3c and Supplementary Figure 2b).

**Figure 3.**
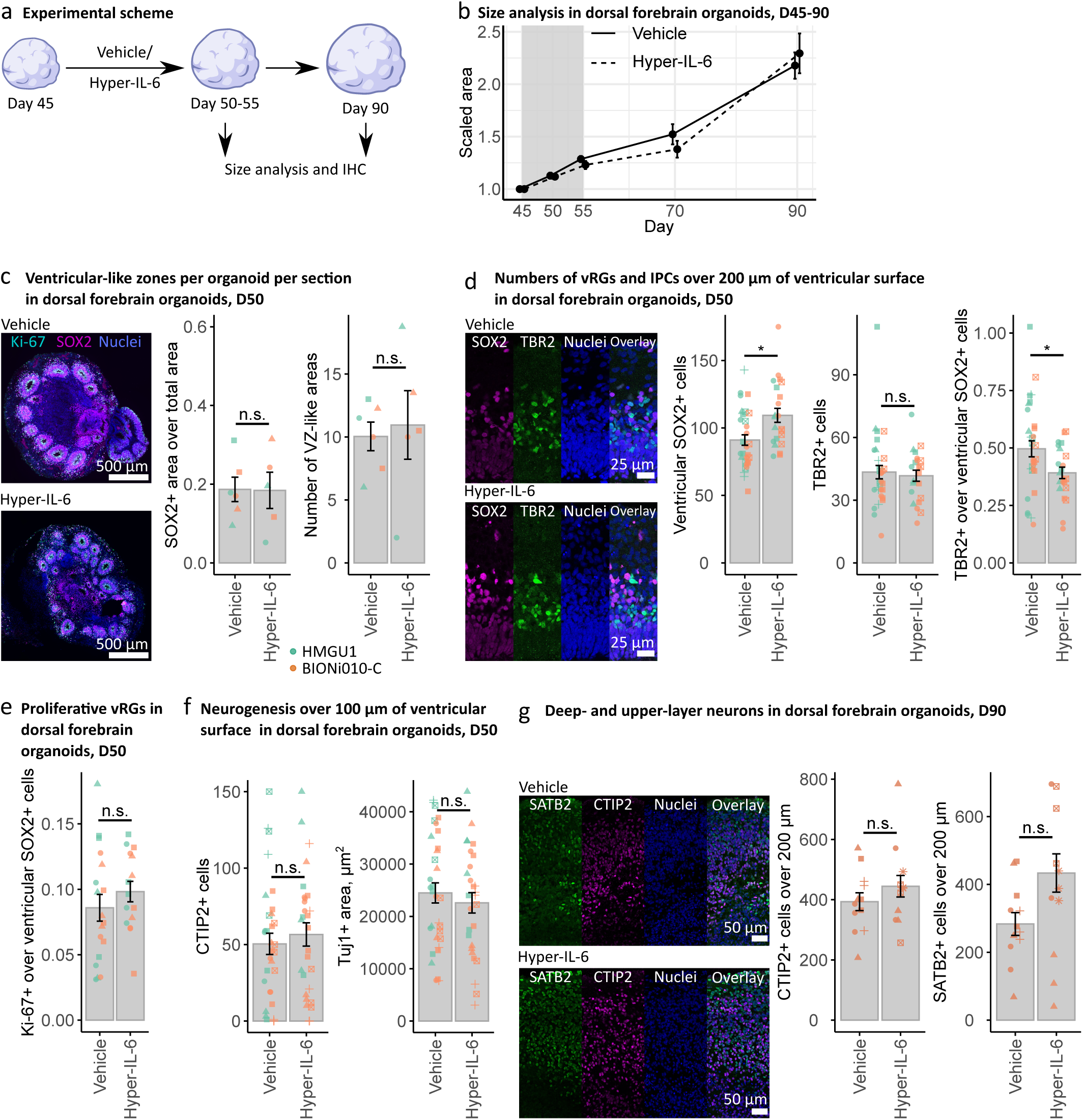
Hyper-IL-6 treatment results in transient changes in cell type composition in DFOs by day 50 of differentiation. a, Experimental timeline and sample harvesting scheme. b, Hyper-IL-6 treatment does not lead to changes in size of the DFOs during the treatment (days 45-55, grey area) and at least up to day 90 of differentiation. Vehicle (n = 5-32) and Hyper-IL-6 (n = 5-32) DFOs from one batch per iPSC line, two iPSC lines. c, Hyper-IL-6 treatment does not lead to changes in the proportion of the SOX2-positive areas and the number of the VZ-like regions in the DFOs. Ki-67 (cyan), SOX2 (magenta), Nuclei stained by DR (blue). Four sections of each DFO were analyzed. Each dot represents the mean value of four sections in a single organoid. Color represents cell line of origin, HMGU1 (green), BIONi010-C (orange). Vehicle (n = 6) and Hyper-IL-6 (n = 5) DFOs from one batch per iPSC line. d, Hyper-IL-6 treatment leads to an increase in SOX2-positive radial glia cells. Hyper-IL-6 treatment does not change the number of TBR2-positive intermediate progenitor cells. Hyper-IL-6 treatment leads to decreased proportion of TBR2-positive over ventricular SOX2-positive cells. Each dot represents individual VZ/SVZ-like region, 2-6 regions imaged per organoid. Color represents cell line of origin, HMGU1 (green), BIONi010-C (orange). Shape corresponds to individual DFO. Vehicle (n = 8) and Hyper-IL-6 (n = 6) DFOs from one batch per iPSC line. e, Hyper-IL-6 treatment does not lead to changes in proportion of Ki-67-positive proliferative cells over SOX2- positive cells. Each dot represents individual VZ-like region, 2-3 regions imaged. Color represents cell line of origin, HMGU1 (green), BIONi010-C (orange). Shape corresponds to individual DFO. Vehicle (n = 6) and Hyper-IL-6 (n = 5) DFOs from one batch per iPSC line. f, Hyper-IL-6 treatment does not lead to changes in CTIP2-positive deep-layer excitatory neurons as well as in the Tuj1-positive area. Each dot represents individual region of interest with VZ-like zone, 2-5 regions imaged per organoid. Color represents cell line of origin, HMGU1 (green), BIONi010-C (orange). Shape corresponds to individual DFO. Vehicle (n = 10) and Hyper-IL-6 (n = 8) DFOs from one batch per iPSC line. g, Hyper-IL-6 treatment does not change the number of CTIP2-positive deep-layer and SATB2-positive upper-layer excitatory neurons at day 90 of differentiation. Each dot represents individual region-of-interest representing a putative CP-like region, 2-3 regions per DFO imaged. All organoids generated from BIONi010-C iPSC line. Shape corresponds to individual DFO. Vehicle (n = 4) and Hyper-IL-6 (n = 5) DFOs from two batches. In all panels, bars represent mean, error bars represent ± SEM. Comparisons were analyzed using Aligned Rank Transform (ART) ANOVA: n.s., non-significant p-value > 0.05, *, p value < 0.05.

The number of SOX2-positive cells within individual VZ-like regions was higher in Hyper-IL-6-exposed organoids (Figure 3d) while the number of TBR2-positive IPCs remained stable upon exposure to Hyper-IL-6 (Figure 3d). The proportion of TBR2-positive IPCs over SOX2-positive RGs was decreased (Figure 3d). The proliferation rate in vRGs, measured as the proportion of Ki-67-positive cells to SOX2+ cell count, was not altered at both ages of the organoids (Figure 3e and Supplementary Figure 3c). We also observed no differences in the count of CTIP2-positive deep-layer neurons as well as the total Tuj1-positive neuron-enriched area upon Hyper-IL-6 exposure at day 50 of differentiation (Figure 3g). Together, these data suggest minor changes in the counts of SOX2-positive RGs, which do not result in an overproduction of IPCs or neurons within 10 days. We also investigated the long-term effect of the Hyper-IL-6 treatment by analyzing the number of CTIP2- and SATB2-positive deep and upper layer neurons at day 90 of differentiation, which represents a more mature stage of human neocortical development when upper layer neurons are present (Figure 3g). We found no significant changes in both CTIP2- and SATB2-positive cell counts (Figure 3g). Coherent with the absence of clinical observations of the gross anatomical malformations in neocortical development upon MIA, we show that Hyper-IL-6 treatment can elicit a functional response in DFOs but no overt cytoarchitectural changes after 6-7 weeks in culture.

### Human dorsal forebrain organoids recapitulate the cellular composition of human neocortex at midgestation

Taken together, our transcriptomic and immunohistochemical analysis indicated that there might be cell-type specific responses to Hyper-IL-6 treatment with strongest effects expected in RGs. To investigate whether Hyper-IL-6 exposure results in cell type-specific molecular changes, we treated DFOs using our established paradigm and profiled individual organoids using scRNA-seq at day 53 of differentiation (Figure 4a). We profiled two organoids per experimental condition originating from two male iPSCs lines. After quality and doublet filtering (quality metrics provided in Extended Data Table 1 and Supplementary Figure 4a), we analyzed the transcriptomes of 11 162 cells, recovering an average of 3 200 unique transcripts per cell. Using Louvain clustering, we identified 16 transcriptionally distinct clusters (Figure 4b). A combination of DGE analysis and analysis of common cell type marker expression allowed us to characterize all clusters. Using this approach, we identified multiple types of progenitor cells and neurons as well as cell states, including proliferative and metabolic status (Figure 4c and Supplementary Figure 4b). Importantly, our cell type classifications were in broad agreement with previous scRNAseq data from DFOs and primary neocortical single-cell transcriptomes (35, 67, 68) (Supplementary Figure 4c). Next, we assessed the neocortical identity of different clusters using VoxHunt (51). First, we used the E13 Allen Developing Mouse Brain Atlas dataset as a reference for the expression of regional identity markers (69). Next, these markers were employed to correlate transcriptomes of our cell clusters with human neocortical transcriptomes from BrainSpan (70). This analysis revealed a subset of neuronal clusters whose regional marker expression did not correlate with neocortical markers (Figure 4d and Supplementary Figure 4d). Cells originating from these non-neocortical neuronal clusters were omitted from further analysis resulting in 8 130 cells with dorsal and medial forebrain identity constituting the final dataset. We reclustered the cells in the final dataset for downstream analysis and obtained 11 clusters (Figure 4e). Analysis of cell type proportions within individual organoids did not reveal major shifts in cellular composition upon Hyper-IL-6 treatment (Figure 4f and Supplementary Figure 4e).

**Figure 4.**
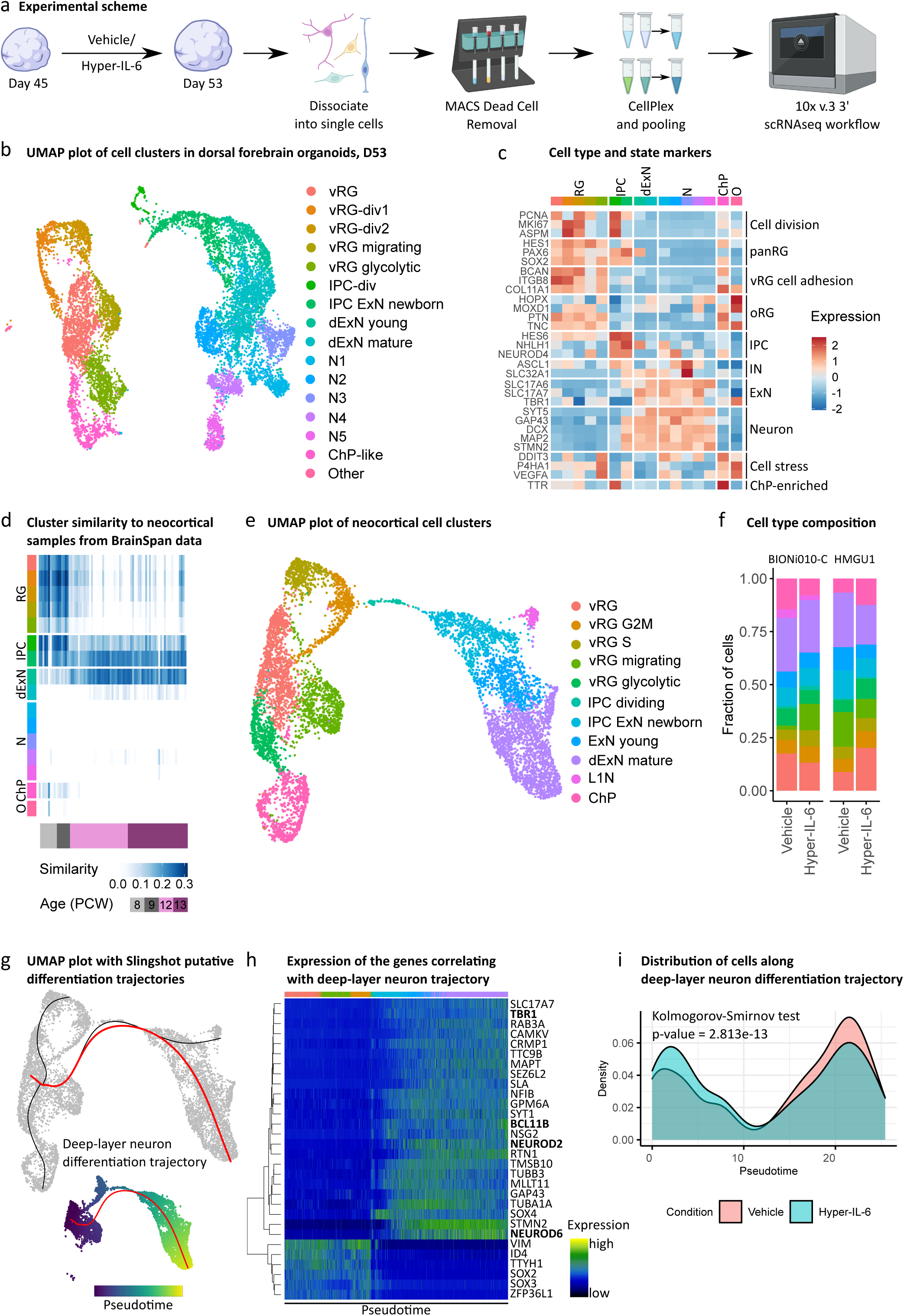
Single-cell RNA sequencing of the dorsal forebrain organoids at day 53 of differentiation upon Hyper-IL-6 and Vehicle treatment. a, Schematic overview of the experimental design, including dissociation of dorsal forebrain organoids, dead cell removal and sample multiplexing by CellPlex. b, UMAP with 16 clusters with cell types labeled by color. Data from n = 2 organoids per condition. vRG, ventricular radial glia; vRG-div, dividing ventricular radial glia; IPC, intermediate progenitor cell; dExN, deep-layer excitatory neuron; N, neuron; ChP, choroid plexus. c, Heatmap of gene expression level for selected known markers of neural cell types across clusters. Colors in the upper bar represent cell types from b. oRG, outer radial glia; IN, inhibitory neuron. d, Heatmap of similarity metric of VoxHunt algorithm comparing organoid clusters with human neocortical RNAseq data from BrainSpan using brain regional markers obtained from Mouse Brain Atlas at E13. Colors in the left bar represent cell types from b. Data from n = 4 organoids. e, UMAP with 11 clusters resulting from subclustering of cells with dorsal and medial forebrain regional identity. f, Bar plots of the proportions of cell types by organoid in two cell lines. g, Putative differentiation trajectories (generated by Slingshot) overlaid over UMAP, deep-layer excitatory neuron differentiation trajectory in red. h, Heatmap of gene expression of genes correlating with progression through pseudotime over deep-layer neuron differentiation trajectory as assessed by random forest classifier. Colors in the bar correspond to cell types from e. Hierarchical clustering of the genes shown on the left. Data from n = 4 organoids. i, Density plot of cells over deep-layer neuron differentiation trajectory by experimental condition. Kolmogorov-Smirnov test. Data from n = 2 organoids per condition.

However, the reconstruction of lineage trajectory showed a shift towards earlier pseudoage in Hyper-IL-6-treated organoids (Figure 4g-i). Together with immunohistochemical data, this result suggests a minor increase of the RGs upon Hyper-IL-6 treatment.

### Differential gene expression analysis suggests disturbed protein translation in radial glial cells

We aimed to characterize cell type-specific transcriptional alterations upon Hyper-IL-6 treatment by performing DGE analysis. In order to focus on cell types rather than cell states, we first unified clusters of vRGs belonging to different stages of the cell cycle into a “cycling vRG” metacluster (Figure 5a). We identified a higher number of DEGs in the cycling vRG cells compared to all other cell clusters (Figure 5b,c). Among the upregulated DEGs in the cycling vRG cells, we identified transcription factors (TFs) STAT3, NR2F1, and NR2F2, among others, based on the comparison with the human TFs list (53) (Figure 5c). Gene set enrichment analysis (GSEA) of DEGs suggested innate immune response-related genes as well as genes involved in the regulation of cell proliferation and gliogenesis being upregulated. This result agrees with previous reports showing that JAK/STAT pathway activation is pro-proliferative and -gliogenic in NPCs (62). Among the gene ontology (GO) terms enriched for downregulated DEGs in cycling vRGs, we found translation initiation and protein targeting to ER (Figure 3d). We also analyzed module expression of the genes based on the significantly de-regulated GO terms identified by Kalish and colleagues (16). We found significant downregulation of translation initiation and cytoplasmic translation in cycling vRG while cytoplasmic translation was upregulated in deep-layer excitatory neurons (Figure 5e). Downregulation of the protein translation-related genes in cycling vRG is in agreement with findings from rodent studies reporting widespread disruption of protein translation in pups’ brains upon MIA exposure (Supplementary Figure 5a,b) (16, 64). Neuronal downregulation of protein translation was suggested to be causal for mediating behavioral abnormalities (16). Here, we provide evidence that abnormal protein translation is evident already on the level of RGs preceding potential abnormalities in the neurons.

**Figure 5.**
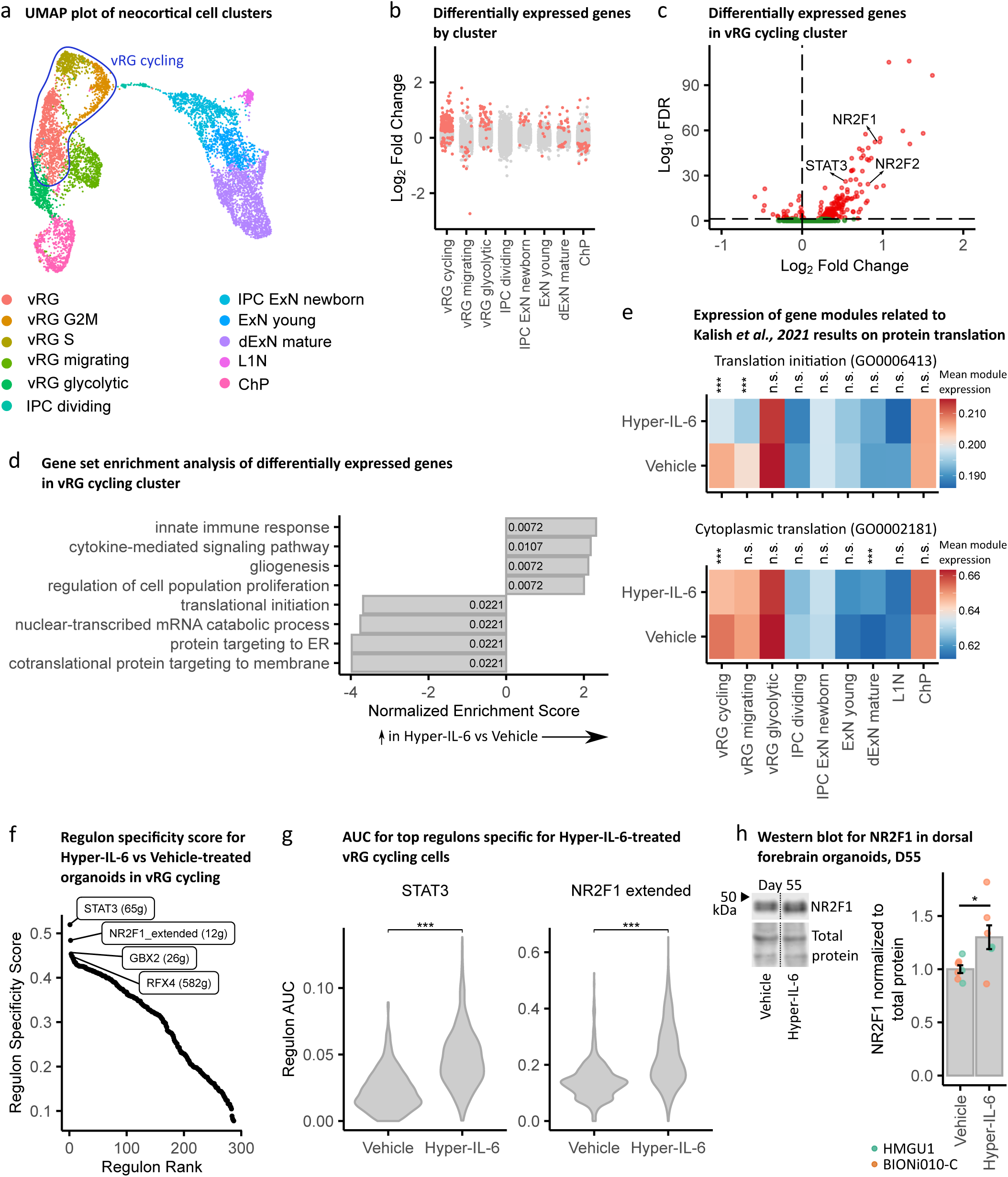
Single-cell DGE and transcriptional networks change upon Hyper-IL-6 treatment. a, vRG, vRG S and vRG G2M clusters unified into cycling vRG metacluster in the UMAP with clusters with cell types resembling forebrain regional identity from Figure 4e. b, Strip plot displaying DEGs between Hyper-IL-6 and Vehicle-treated organoids in red (FDR < 0.05). The x axis displays cell clusters from a. L1N is missing because these cells were not present in all samples. c, Volcano plot of Hyper-IL-6- dependent gene expression in cycling vRG metacluster. Red dots indicate statistical significance (FDR < 0.05). Positive log2 Fold Change indicates higher gene expression in Hyper-IL-6-treated relative to Vehicle-treated cells. d, Gene set enrichment analysis of differentially upregulated genes (FDR < 0.05) between Hyper-IL-6 and Vehicle-treated cycling vRG cells from c. The x axis displays normalized enrichment score. Numbers inside the bars represent adjusted p-values for differential enrichment. e, Heatmap representing mean module expression assessed by UCell by treatment condition across cell clusters from a. Comparisons were analyzed using t-test, Bonferroni adjusted p-values: ***, p- value < 0.001. f, Rank for regulons in Hyper-IL-6-treated cycling vRG cells based on regulon specificity score (RSS) found by SCENIC. g, Regulon area under the curve (AUC) for STAT3 and NR2F1-driven regulons in cycling vRG cells between conditions. Comparisons were analyzed using t-test: ***, p- value < 0.001. h, Representative Western Blot image and quantification measuring NR2F1 (top) normalized to Total Protein Stain (bottom) in dorsal forebrain organoids at day 55 of differentiation. In the plot, each dot represents an individual organoid. Color represents cell line of origin, HMGU1 (green), BIONi010-C (orange). Vehicle (n = 7) and Hyper-IL-6 (n = 7) DFOs from one batch per iPSC line. Bars represent mean, error bars represent ± SEM. Comparisons were analyzed using Aligned Rank Transform (ART) ANOVA: *, p-value < 0.05.

To analyze relevance of the DEGs for ASD phenotype we performed two types of analysis. First, gene module expression analysis of the ASD-relevant gene groups in our single-cell dataset revealed significant deregulation of gene groups “Cytoskeleton” and “Neuronal communication” specifically in RGs based on gene sets reported by Satterstrom and colleagues (Supplementary Figure 5c) (71). However, SFARI genes of categories 1-4 by the old classification and categories 1-2 were not differentially enriched among DEGs in our single-cell clusters by the new classification (Supplementary Figure 5d).

We further performed SCENIC analysis to better characterize gene networks deregulated by Hyper-IL-6 treatment specifically in vRG cycling cells (56). The highest specificity scores (58) were obtained by STAT3 and NR2F1-driven regulons (Figure 5f). The activity of these regulons was significantly higher in Hyper-IL-6-treated cells (Figure 5g). While increased activity of STAT3-mediated gene expression further validates our model (Supplementary Figure 5e) in line with RNAseq and Western Blot data, NR2F1 has not been reported as differentially expressed or active in rodent models of MIA (16, 64). Gene set overrepresentation analysis of the genes in NR2F1 regulon revealed genes involved in synapse organization and cell migration (Supplementary Figure 5f,g). We confirmed the upregulation of NR2F1 on the protein level in Hyper-IL-6-treated organoids at day 55 of differentiation (Figure 5h). Additionally, RNAseq data also showed an upregulation of NR2F1 in Hyper-IL-6-treated organoids at day 55 of differentiation (log2 Fold Change = 0.705495957, adjusted p-value = 0.11213299).

## Discussion

In this study, we close the gap between human epidemiological studies and mechanistic studies in animal models by establishing a human 3D model to study the effects of MIA on the developing brain using iPSCs-derived brain organoids and cytokine signaling activation.

### IL-6 as a molecular mediator of MIA

Epidemiological studies, as well as animal and 2D *in vitro* models of MIA, have revealed possible molecular mediators of MIA, including IL-6, IL-17α, TNF-α, and IFN-γ, that drive the adverse neurodevelopmental outcomes (11, 16, 17, 22, 72). Among those, there is strong evidence for IL-6 playing a key role (11, 18). In our model, we therefore chose to focus on IL-6 signaling. In order to show the validity of this approach, we confirmed that the molecular machinery required for responding to IL-6 is expressed in the developing human neocortex with a similar expression pattern in DFOs. Since only microglia express IL6R in the developing human brain (32), we hypothesized that neuroectodermal cells can only be activated by IL-6, if IL6R is provided in *trans* from microglia. We model *trans* signaling in DFOs devoid of microglia by applying Hyper-IL-6, a chimeric protein consisting of the soluble IL6R and IL-6 bound through an oligopeptide link (73). We observed that only Hyper-IL-6, not IL-6 itself, activates JAK/STAT signaling validating our hypothesis (Figure 2). Therefore, we chose to model MIA with Hyper-IL-6 in DFOs devoid of microglia. Similarly, we suggest that IL-17α may also mediate neurodevelopmental outcomes following MIA (16, 17) through microglia, since transcriptomics data suggests that IL17RA is expressed predominantly in microglia and only sparsely in cells of neuroectodermal origin (32, 33). In conclusion, our model can be expanded on in the future in DFOs with microglia (74–76) recapitulating the interaction of radial glia and microglia in MIA in the developing rodent neocortex (77, 78).

### Cellular effects of MIA on neocortical development: modulation of neurogenesis

It has been shown that MIA during the first and early second trimester poses the highest risk for developing MIA-associated neurodevelopmental deficits (5). We therefore focus on an early time point in organoid differentiation, exposing organoids from Day 45 to Day 55 with Hyper-IL-6 modeling the period of early midgestation (32). Infections usually last several days, prompting us to treat organoids for 5 to 10 days with Hyper-IL-6. Immunohistochemical analysis of different cell populations showed increased numbers of Sox2-positive vRGs, while not showing any effect on IPCs and neurons after five days of treatment (Figure 3). Similarly, the reconstruction of the deep-layer neuron differentiation trajectory from our scRNAseq dataset revealed a higher proportion of progenitor cell populations compared to neuronal cell populations upon Hyper-IL-6 exposure (Figure 4). These results partially recapitulate the results from a mouse model of MIA where poly(I:C) treatment resulted in higher numbers and proliferative capacity of PAX6-positive vRGs (18). In the mouse, however, MIA also led to an overproduction of deep layer neurons (18). When we quantified neuronal populations on Day 90, 35 days after the end of Hyper-IL-6 treatment, we did not find any significant changes in long-term neuronal output (Figure 3). This discrepancy may be due to species-specific differences in the cellular responses to MIA or differences in the quality of MIA induced by poly(I:C) and Hyper-IL-6 treatment respectively. Taken together, we model the effect of MIA on the early phase of human neocortical neurogenesis. We expect that future studies will use our model to reveal how MIA affects later neurodevelopmental processes in a human context, focusing for example on the effect of MIA on outer or basal radial glia cells, which are abundant in the human but almost absent from the mouse neocortex (79–81).

### Molecular effects of MIA on neocortical development: changes gene expression associated with immune response, protein translation, and NR2F1 signaling

In order to determine which molecular changes are induced in our MIA model and how they compare mechanistically to previous findings in other models, we performed transcriptomic analysis. In RNAseq data, the top upregulated genes were at the interface of innate and adaptive immune responses: MHCI and B2M, which encodes the ancillary protein necessary for the MHCI assembly (82). Analogously, Warre-Cornish and colleagues also found expression of MHCI proteins in iPSC-derived neural precursor cells treated with IFN-γ, a major modulator of the innate immune response (22). This resulted in the abnormal neurite growth upon neuronal differentiation of IFN-γ- treated NPCs which was rescued by a B2M knockdown (22). The authors suggest that altered neurite growth may be related to similar observations made in iPSC-derived neurons from patients with ASD (83–85). In our study, WGCNA analysis showed co-expression of MHCI genes with RG-enriched genes (Figure 2) suggesting a similar mechanism of IL-6 action as found for IFN-γ-treated NPCs.

Upregulation of MHCI proteins was also reported in a mouse model of MIA (86) and HLA-A alleles are differentially associated with ASD in humans (87). Therefore, our data together with previous results further strengthens the link between the cellular effects of MIA and altered neurodevelopmental processes observed in ASD (22, 86).

Since our WGCNA analysis indicated that RGs might be specifically affected by Hyper-IL-6 treatment, we performed cell type-specific differential gene expression analysis by performing single-cell RNAseq. Indeed, we found the highest number of differentially expressed genes in the vRG cells.

Among the downregulated genes were many involved in protein translation, consistent with the results obtained from diverse cell types including radial glia in a mouse model of MIA (16). In contrast to the data provided by Kalish and colleagues, our data identifies deregulation not only of cytoplasmic translation but also protein targeting to the endoplasmic reticulum (ER) and plasma membrane. Changes in protein secretion may be biologically relevant especially in the human developing neocortex, which has a different extracellular matrix composition compared to rodents (31). Notably, it has previously been reported that ER stress may protract developmental shift from direct to indirect neurogenesis (88). The finding that ER-related processes are dysregulated in our model may explain the decreased proportion of TBR2-positive IPCs to SOX2-positive vRGs revealed by immunohistochemical analysis. Finally, downregulation of translation machinery components may alter the translation landscape in a cell type-specific manner (89, 90). In the context of new findings regarding the role of translational regulation on cell fate acquisition in murine RGs (89, 90), investigating cell type-specific alterations in protein translation may be possible in the future with current developments for example in single-cell (91) and ultrasensitive (92) Ribo-seq.

Since gene expression has correlative structure where expression of groups of genes is driven by binding of TFs to the respective genomic loci, in so-called regulons, these regulons can be perceived as functional entities of gene expression (56). Analysis of regulon activity in cycling vRGs revealed two major TFs driving transcriptional changes in Hyper-IL-6-treated organoids, STAT3 and NR2F1 (Figure 5). STAT3 validates our model, since STAT3 is the known major transcription factor downstream of IL-6/JAK/STAT signaling (62). NR2F1 has not been reported to be involved in mediating the effects of MIA but it is an important TF with multiple cell type- and cortical/brain region-specific functions (36, 93). In dorsal forebrain RGs, it determines the balance between proliferation and differentiation by acting in counterbalance with PAX6 (94). Given that in comparison with murine data we do not observe neuronal overproduction in response to MIA, we propose the hypothesis that NR2F1 is upregulated in RGs as a protective factor against the neotenic effect of IL-6 and that this effect may differentiate mouse and human model systems.

Taken together, our findings corroborate the findings of Kalish and colleagues who point towards a causal role of abnormalities in protein translation for development of ASD-like phenotype upon MIA exposure (16). Additionally, the upregulation of genes involved in the innate immune response, including those of MHCI, partially recapitulates the effects of IFN-γ exposure in NPCs (22). Therefore, our model demonstrates the utility of human brain organoids for investigating the molecular mechanisms underlying the effects of MIA on human neurodevelopment. While our study focused only on male DFOs, future studies may determine whether similar effects are seen in female DFOs or whether female DFOs possess inherent resilience mechanisms as suggested by mouse models (16).

In conclusion, our study established a model of MIA in brain region-specific organoids and provides a framework for future studies of the effects of MIA on multiple organizational levels, from subcellular to tissue-wide. Our results corroborate previous results from the rodent and 2D *in vitro* model systems (16, 18, 22, 64). However, we also extend our knowledge of cell types primarily affected by MIA, by showing that radial glia cells are specifically vulnerable to IL-6 signaling, shifting them towards a younger pseudoage along the neural differentiation trajectory. Finally, the brain organoid model of MIA may serve as a platform for investigating the effects of the interaction of genetic and environmental mediators of complex neurodevelopmental diseases such as ASD (59, 95, 96).

### Data availability

Sequencing data generated in this study will be accessible at the repository Array Express under accession number ().

## Acknowledgements

We thank Elisabeth Gustafsson, Lea Fischer, Jasmin Treu, Maximilian Feige, Christina Kulka, and Felix Hildebrand for technical support and NCCT for scRNA-sequencing support. We thank Dmitry Kobak, Philipp Berens, and Peter Heutink for advice on single-cell transcriptomics data analysis. We thank Arnold Kriegstein and his team for providing primary tissue samples. We thank the Helmholtz Zentrum München Deutsches Forschungszentrum für Gesundheit und Umwelt for providing HMGU1 cells and EBiSC Bank for providing BIONi010-C. The EBiSC Bank acknowledges Bioneer as the source of the human induced pluripotent cell line BIONi010-C (97). We thank John Blair and Helen Beteup for advice on protein extraction from brain organoids. We thank Andrii Zadaianchuk for reviewing scripts generated for single-cell transcriptomics data analysis. We thank Hansjürgen Volkmer and Claudia Buss for critical feedback on the manuscript.

## Contributions

KS designed the study, performed experiments, performed data analysis, statistical analysis, and prepared the manuscript; TK, SK, EA, ZY and KB performed experiments and revised the manuscript; SM conceived and designed the study, supervised the work, and prepared the manuscript.

