## Supplementary figures for "Human brain organoid model of maternal immune activation identifies radial glia cells as selectively vulnerable"

### Supplementary Figure 1

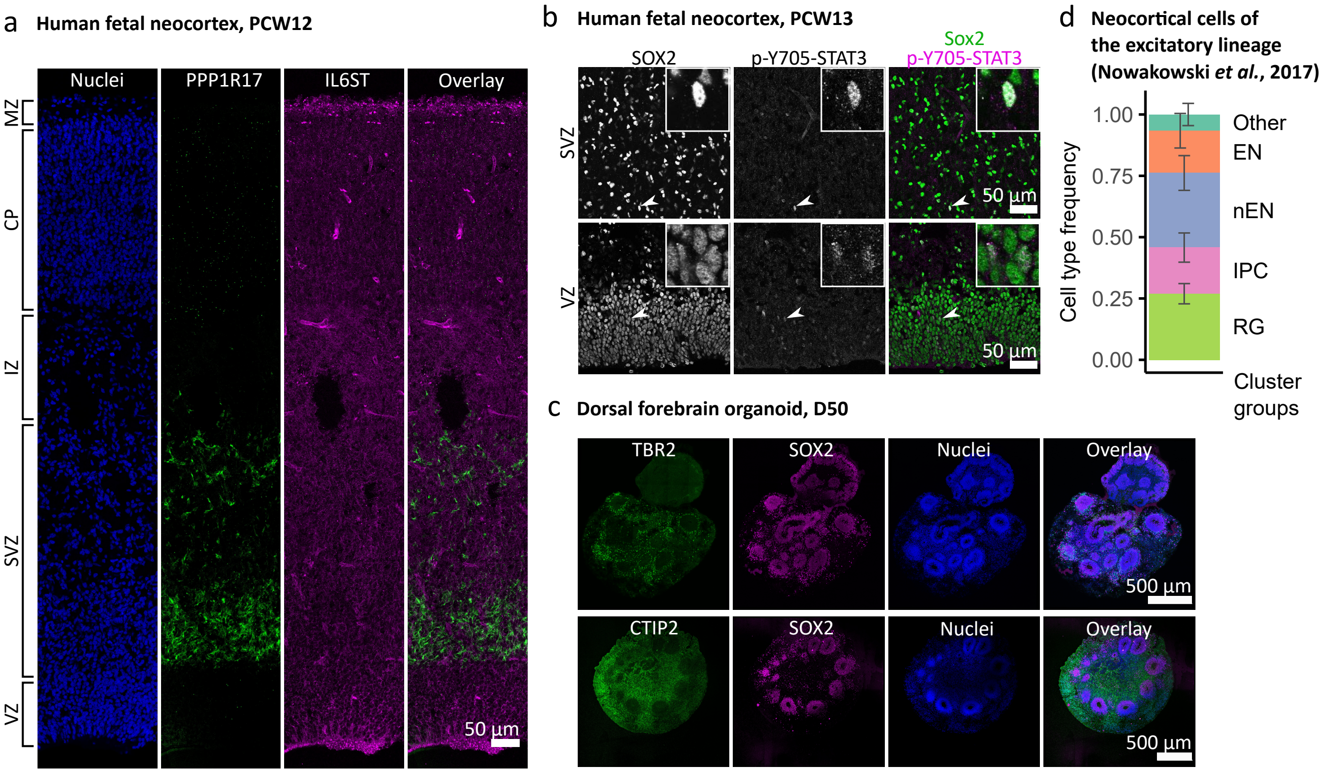

**Supplementary Figure 1. Related to Figure 1. Characterization of IL-6 cascade in human neocortex and dorsal forebrain organoids.** a, Confocal tilescan image across human neocortex at PCW12 immunostained for IL6ST (magenta) and PPP1R17 (green). b, p-Y705-STAT3 immunofluorescence in human neocortex at PCW13 shows co-localization with the marker of radial glia cells, SOX2. c, At day 50 of differentiation, dorsal forebrain organoids consist preferentially of SOX2-positive radial glia, TBR2-positive intermediate progenitor cells and CTIP2-positive deep-layer excitatory neurons. d, Excitatory lineage of the human neocortical cells at PCW9-15 largely consists of the radial glia, intermediate progenitor cells, newborn and mature neurons in line with organoid cell type composition based on single-cell transcriptomes from Nowakowski *et al, 2017*.

### Supplementary Figure 2

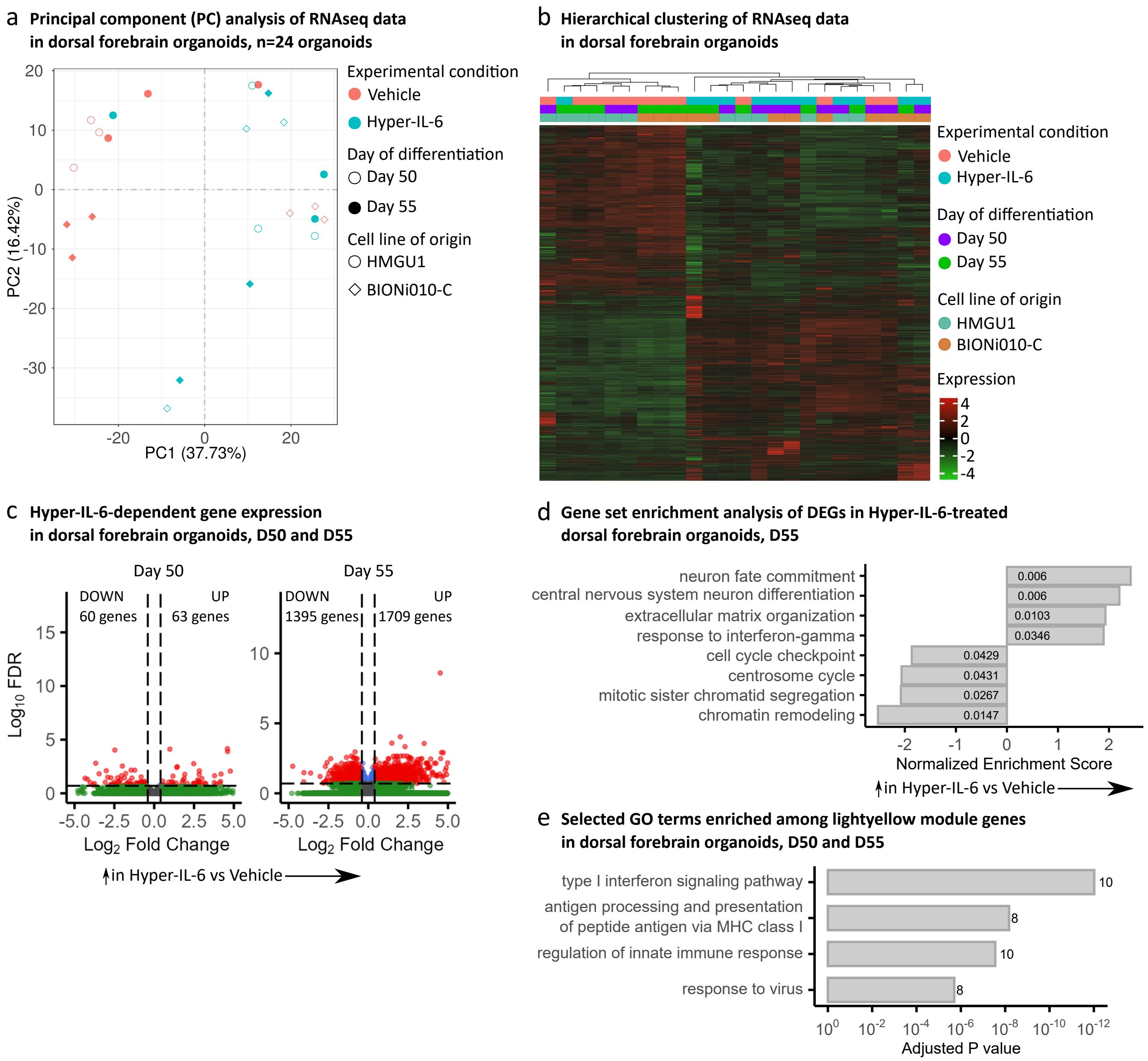

**Supplementary Figure 2. Related to Figure 2. Quality control of RNAseq data.** a, Principal component analysis of RNAseq data from n=24 organoids from 2 iPSCs cell lines (BIONi010-C and HMGU1), two time points over organoids differentiation (D50 and D55) and two experimental conditions (Vehicle and Hyper-IL-6). b, Hierarchical clustering of RNAseq data showing clustering of samples on the basis of experimental condition, day of differentiation and iPSC line. Clustering was performed based on all expressed genes. c, Volcano plot of Hyper-IL-6-dependent gene expression in dorsal forebrain organoids at days 50 and 55 of differentiation treated for 5 and 10 days, respectively. Red dots indicate statistical significance (FDR < 0.2, absolute log2 Fold Change > 0.4). Positive log2 Fold Change indicates higher gene expression in Hyper-IL-6-treated relative to Vehicle-treated dorsal forebrain organoids. Data from n=6 organoids per condition per day of differentiation (two cell lines). d, Gene set enrichment analysis of differentially upregulated genes (FDR < 0.2, absolute log2 Fold Change > 0.4) between Hyper-IL-6 and Vehicle-treated organoids at day 55 of differentiation. The x axis displays normalized enrichment score. Numbers inside the bars represent adjusted p-values for differential enrichment. e, GO overrepresentation analysis of genes constituting lightyellow WGCNA module. The x axis displays the adjusted p-value. Numbers next to the bars represent the number of the differentially expressed genes belonging to the GO term. Data from n=12 organoids per condition (two cell lines, day 50 and 55 analyzed together).

### Supplementary Figure 3

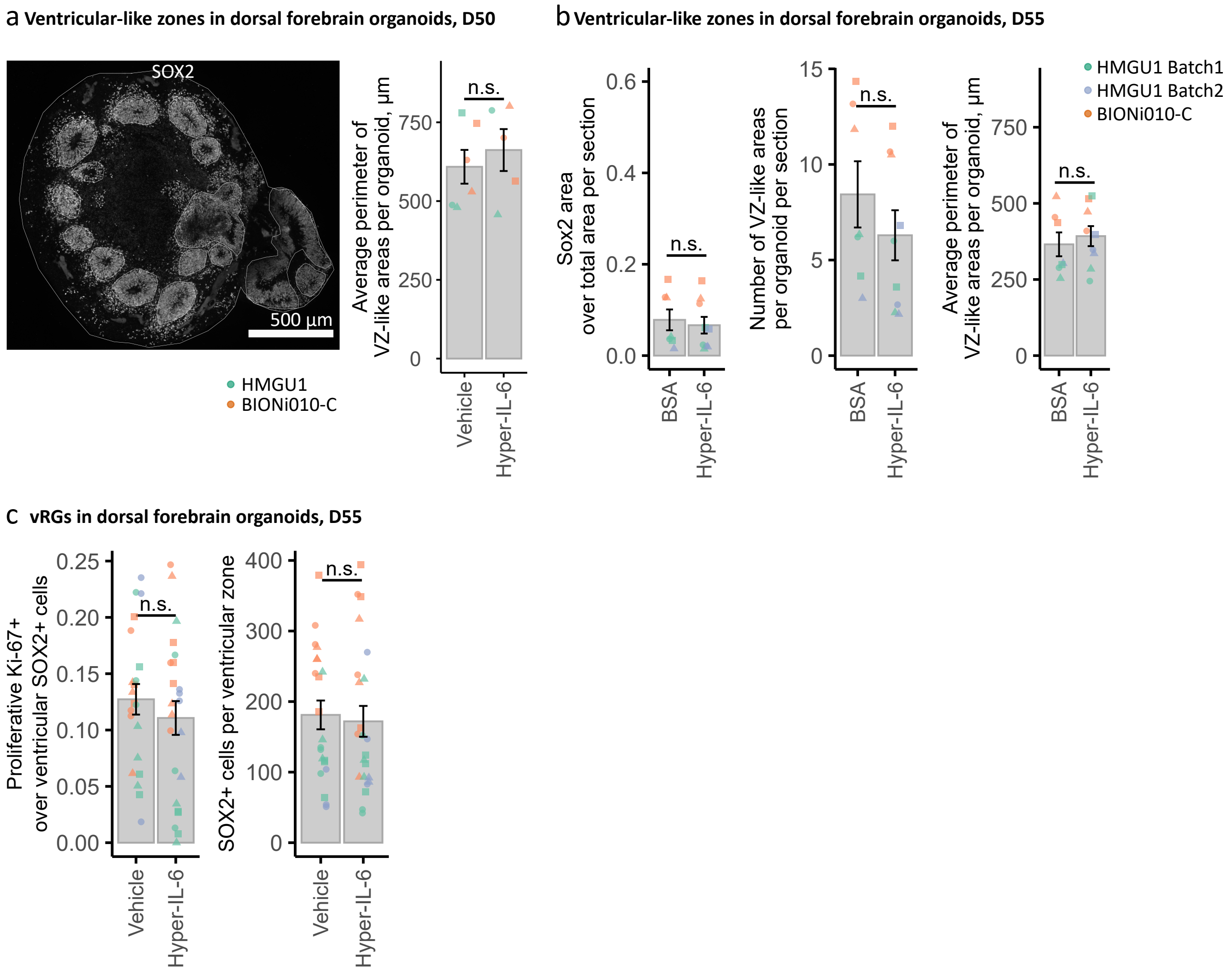

**Supplementary Figure 3. Related to Figure 3. Hyper-IL-6 treatment does not result in changes in cell type composition in DFOs by day 55 of differentiation.** a, Hyper-IL-6 treatment does not lead to changes in the average perimeter of SOX2-positive areas. Four sections of each DFO were analyzed. Representative tilescan image displays an exemplary analysis workflow. White lines outline the organoid and SOX2-positive areas within it. In the plot, each dot represents the mean value of four sections in one single organoid. Color represents cell line of origin, HMGU1 (orange), BIONi010-C (green). Vehicle (n = 6) and Hyper-IL-6 (n = 5) DFOs from one batch per iPSC line. b, Hyper-IL-6 treatment does not lead to changes in the average perimeter of SOX2-positive areas, their number and area at day 55 of organoids differentiation. Six sections of each DFO were analyzed. Each dot represents the mean value of six sections in one single organoid. Color represents cell line of origin, HMGU1 (green), BIONi010-C (orange). Vehicle (n = 7) and Hyper-IL-6 (n = 9) DFOs both iPSC lines. c, Hyper-IL-6 treatment does not lead to changes in proportion of Ki-67-positive proliferative cells over SOX2-positive cells on day 55 of differentiation. Hyper-IL-6 does not lead to an increase in SOX2-positive radial glia cells on day 55 of differentiation. Each dot represents individual VZ-like region, 2-3 regions imaged. Color represents cell line of origin and batch of differentiation, HMGU1 (green and violet), BIONi010-C (green). Shape corresponds to individual DFO. Vehicle (n = 7) and Hyper-IL-6 (n = 8) DFOs from both iPSC lines. In all panels, bars represent mean, error bars represent ± SEM. Comparisons were analyzed using Aligned Rank Transform (ART) ANOVA: n.s., non-significant p-value > 0.05.

### Supplementary Figure 4

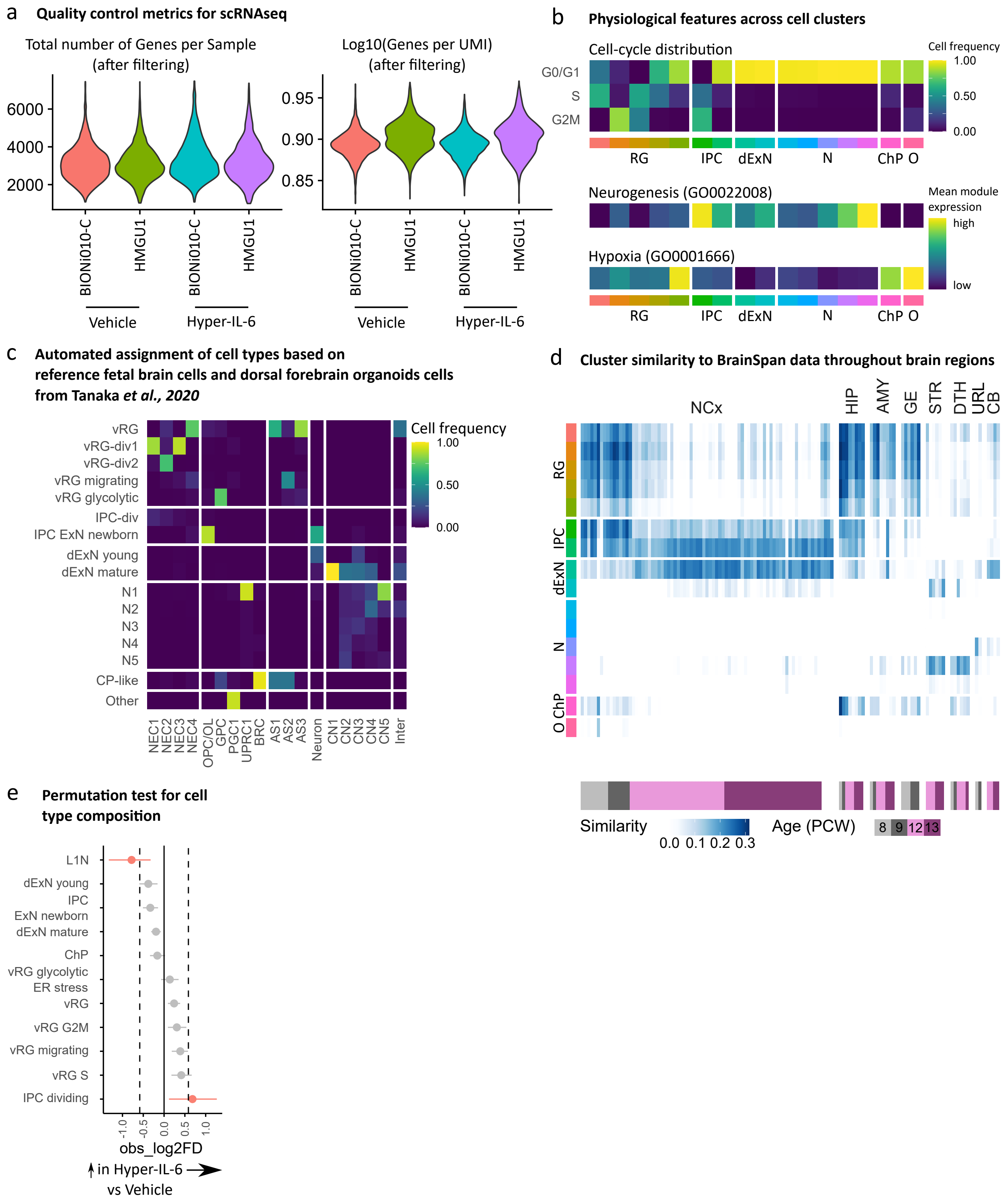

**Supplementary Figure 4. Related to Figure 4. Quality control of scRNAseq and cell type composition analysis.** a, Total number of genes per sample and log10 of genes per UMI per sample in the single cell sequencing dataset. b, Heatmap of physiological feature assessment across clusters. Distribution of the cells in each cluster across cell cycle phases (top), expression of module signature genes for GO terms related to Neurogenesis (GO:0022008, middle) and Hypoxia (GO:0001666, bottom) across cell clusters. Colors in the lower bars represent cell types from Figure 4b. c, Automated assignment of cell cluster identity to the dataset consisting of the human neocortical cells (from Zhong et al., 2018) and dorsal forebrain organoid cells (from Velasco et al., 2019) with cell type labels from Tanaka et al., 2020. Cell frequency is presented column-wise. Cell type labels by Tanaka and colleagues: NEC, neuroepithelial cell; OL, oligodendrocyte; GPC, glia progenitor cell; PGC, proteoglycan-expressing cell; UPRC, unfolded protein response-related cell; BRC, BMP-related cell; AS, astrocyte; CN, cortical excitatory neuron; Inter, interneuron. d, Heatmap of similarity metric of VoxHunt algorithm comparing organoid clusters with human brain RNAseq data from BrainSpan using brain regional markers obtained from Mouse Brain Atlas at E13. Colors in the left bar represent cell types from Figure 4b. e, Permutation test on cell type composition of Hyper-IL-6-treated organoids. Differentially abundant cell types are represented in pink. FDR less than 0.05 and absolute log2 fold change more than 0.58 were considered differentially abundant. In all panels, data from n = 4 organoids from two cell lines (HMGU1 and BIONi010-C) and two experimental conditions (Vehicle and Hyper-IL-6).

### Supplementary Figure 5

a **Venn diagram of differentially downregulated genes in cycling vRGs and RGs from Kalish *et al., 2021***

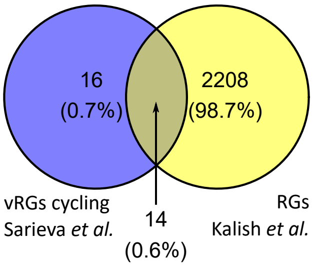


b **Gene set overrepresentation analysis of 14 differentially downregulated genes overlapping in cycling vRGs and RGs from Kalish *et al., 2021***

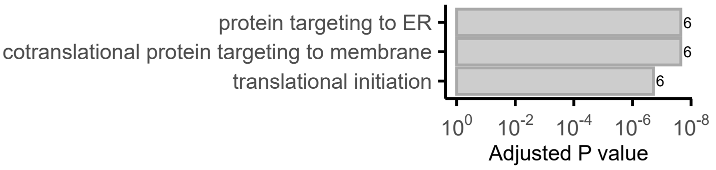


c **Expression of gene modules related to ASD based on Satterstrom *et al., 2020***

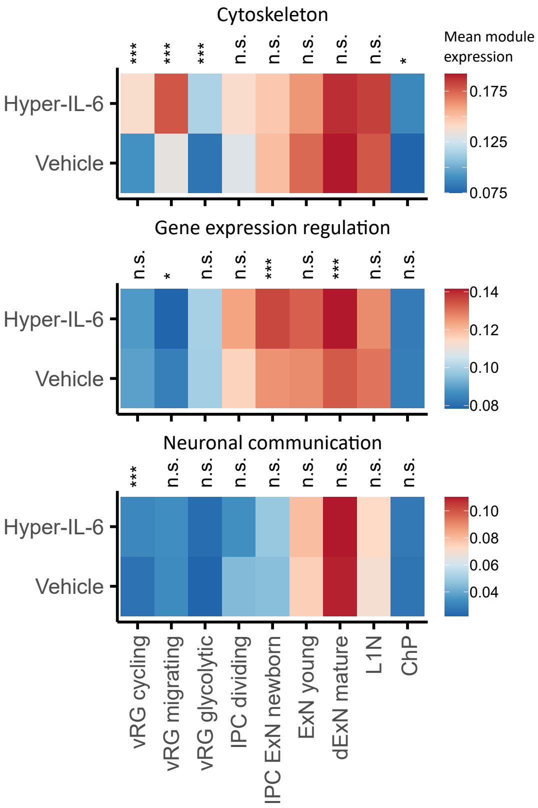


d **Relevance of Hyper-IL-dependent gene expression changes to ASD**

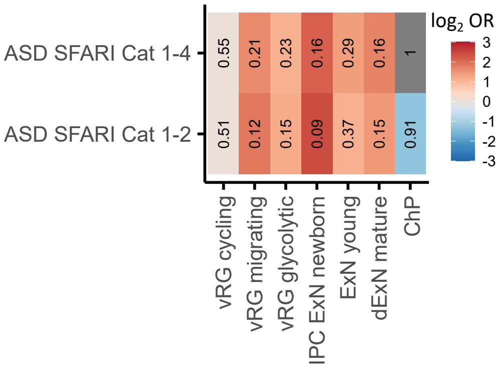


e **Gene set overrepresentation analysis of the genes within STAT3 regulon in cycling vRGs**

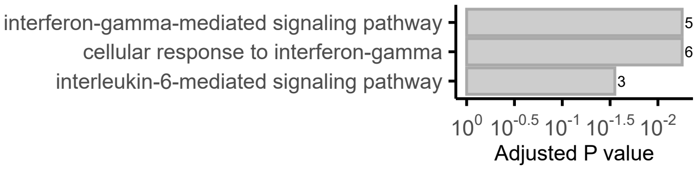


f **Gene set overrepresentation analysis of the genes within NR2F1 regulon in cycling vRGs**

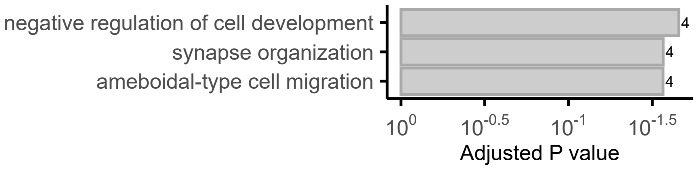


g **Genes within NR2F1 regulon in cycling vRGs**

| Gene name | Protein name |
|---|---|
| ANK2 | ankyrin 2 |
| CLU | clusterin |
| ERBB4 | erb-b2 receptor tyrosine kinase 4 |
| GLIPR1 | GLI pathogenesis related 1 |
| GPC3 | glypican 3 |
| LRRN3 | leucine rich repeat neuronal 3 |
| NFIA | nuclear factor I A |
| NPPC | natriuretic peptide C |
| NR2F1 | nuclear receptor subfamily 2 group F member 1 |
| NR2F2 | nuclear receptor subfamily 2 group F member 2 |
| SEMA5A | semaphorin 5A |
| SPARCL1 | SPARC like 1 |

**Supplementary Figure 5. Related to Figure 5. Single-cell DGE and transcriptional networks changes upon Hyper-IL-6 treatment.** a, Venn diagram showing the intersection of differentially downregulated genes from cycling vRG metacluster and RG cells from Kalish and colleagues. b, GO overrepresentation analysis of overlapping differentially downregulated genes from a. The x axis displays the adjusted p-value. Numbers next to the bars represent the number of the differentially expressed genes belonging to the GO term. c, Heatmap representing mean module expression of ASD-relevant gene groups by treatment condition across cell clusters from Figure 5a. Comparisons were analyzed using t-test, Bonferroni adjusted p-values: n.s., non-significant p-value > 0.05; *, p-value < 0.05; ***, p-value < 0.001. d, Enrichment of ASD risk genes and among DEGs upon Hyper-IL-6 treatment. Log2 odds ratio (OR) is represented by color and adjusted p-values are indicated inside the tiles. Fisher’s exact test; p-values corrected for multiple comparisons by Benjamini-Hochberg method. ASD SFARI Cat 1-4 corresponds to ASD SFARI risk genes from categories 1-4 from SFARI release on 31.10.2019 (old classification); ASD SFARI Cat 1-2 corresponds to ASD SFARI risk genes from categories 1-2 from SFARI release on 20.07.2022 (new classification). e,f, GO overrepresentation analysis of genes belonging to STAT3 (e) and NR2F1_extended (f) regulons as assessed by SCENIC. The x axis displays the adjusted p-value. Numbers next to the bars represent the number of the genes belonging to the GO term. g, Genes belonging to the NR2F1_extended regulon as assessed by SCENIC.
